# A latent variable approach to decoding neural population activity

**DOI:** 10.1101/2020.01.06.896423

**Authors:** Matthew R Whiteway, Bruno Averbeck, Daniel A Butts

## Abstract

Decoding is a powerful approach for measuring the information contained in the activity of neural populations. As a result, decoding analyses are now used across a wide range of model organisms and experimental paradigms. However, typical analyses employ general purpose decoding algorithms that do not explicitly take advantage of the structure of neural variability, which is often low-dimensional and can thus be effectively characterized using latent variables. Here we propose a new decoding framework that exploits the low-dimensional structure of neural population variability by removing correlated variability that is unrelated to the decoded variable, then decoding the resulting denoised activity. We demonstrate the efficacy of this framework using simulated data, where the true upper bounds for decoding performance are known. A linear version of our decoder provides an estimator for the decoded variable that can be more efficient than other commonly used linear estimators such as linear discriminant analysis. In addition, our proposed decoding framework admits a simple extension to nonlinear decoding that compares favorably to standard feed-forward neural networks. By explicitly modeling shared population variability, the success of the resulting linear and nonlinear decoders also offers a new perspective on the relationship between shared variability and information contained in large neural populations.

## 1 Introduction

The increasing ability to record activity from large neural populations now allows for the study of information processing at the level of populations rather than individual neurons [1]. However, the complexity of such recordings also introduces new challenges for data analysis [2, 3]. One popular approach for understanding such complex, high-dimensional data is to search for lower dimensional representations using unsupervised latent variable models [4, 5]. Decoding is an alternative, supervised approach that relates neural activity to variables such as stimuli or behavior. This approach is useful when one is interested in the neural representation of specific variables, and can provide complementary information to related encoding models [6].

Studies across all major neural recording modalities have used decoding analyses to investigate the relationship between neural activity and variables such as external stimuli [7–9], motor outputs [10, 11], decisions [12, 13], and spatial location [10,14,15]. As neural recordings begin to measure simultaneous activity across multiple regions, decoding also promises to play a pivotal role in understanding how information content differs across brain regions [16].

The increasing numbers of neurons that can be simultaneously recorded can make decoding analyses more relevant, but also present challenges to the effective decoding of a given variable from neural activity. Most decoding algorithms show good statistical properties only when the number of observations (e.g. trials) is much larger than the number of features (e.g. neurons) [17]. Current recording technologies focus on increasing the number of neurons, without a concomitant increase in the number of trials, as most experiments are still typically of limited duration (although advances in chronic recordings promise to partially ameliorate this problem [18, 19]). This trend highlights a need to develop new decoding algorithms that are better suited to this many-neuron, few-trial regime.

Here we propose a solution that is tailored to the known structure of neural population activity. Recent studies have demonstrated that variability in neural activity tends to be low-dimensional across many experimental paradigms, and can thus be well-described by latent variable models [20–25]. This suggests that one way of improving a decoding algorithm would be to first estimate low-dimensional variability that is not task-related, remove it from the population responses, and then decode the denoised residual response with reference to the task.

We first provide an intuitive explanation of this decoding framework, and demonstrate the efficacy of a linear version on simulated data where analytic bounds for decoding performance allow a straightforward comparison to other linear decoders such as Linear Discriminant Analysis (LDA) and logistic regression. We then show how this decoding framework can be naturally extended to a nonlinear method through the use of artificial neural networks. Finally, we discuss the implications of these results when analyzing real neural data.

## 2 Results

### 2.1 Describing noise correlations with latent variables

We begin by describing a simple example that elucidates the relationship between latent variables and noise correlations, which will motivate the development of our latent variable decoding framework in the following section. Consider a task in which the subject sees one of two possible stimuli on each trial and must perform a saccade to the presented stimulus, while we record the simultaneous activity of many neurons in a task-relevant brain region. To visualize how the dynamics of neural population activity unfold in time, we perform dimensionality reduction on the trial-averaged responses using principal component analysis (PCA), and project the high-dimensional activity into the first three principal components (Fig. 1A, *bold lines*). Notably, trial averaging is necessary in this case because variability in single-trial population responses from trial-to-trial often introduces many more relevant dimensions into the population response, and may disrupt visualization of the trajectories that only depend on the fixed experimental conditions.

**Figure 1:**
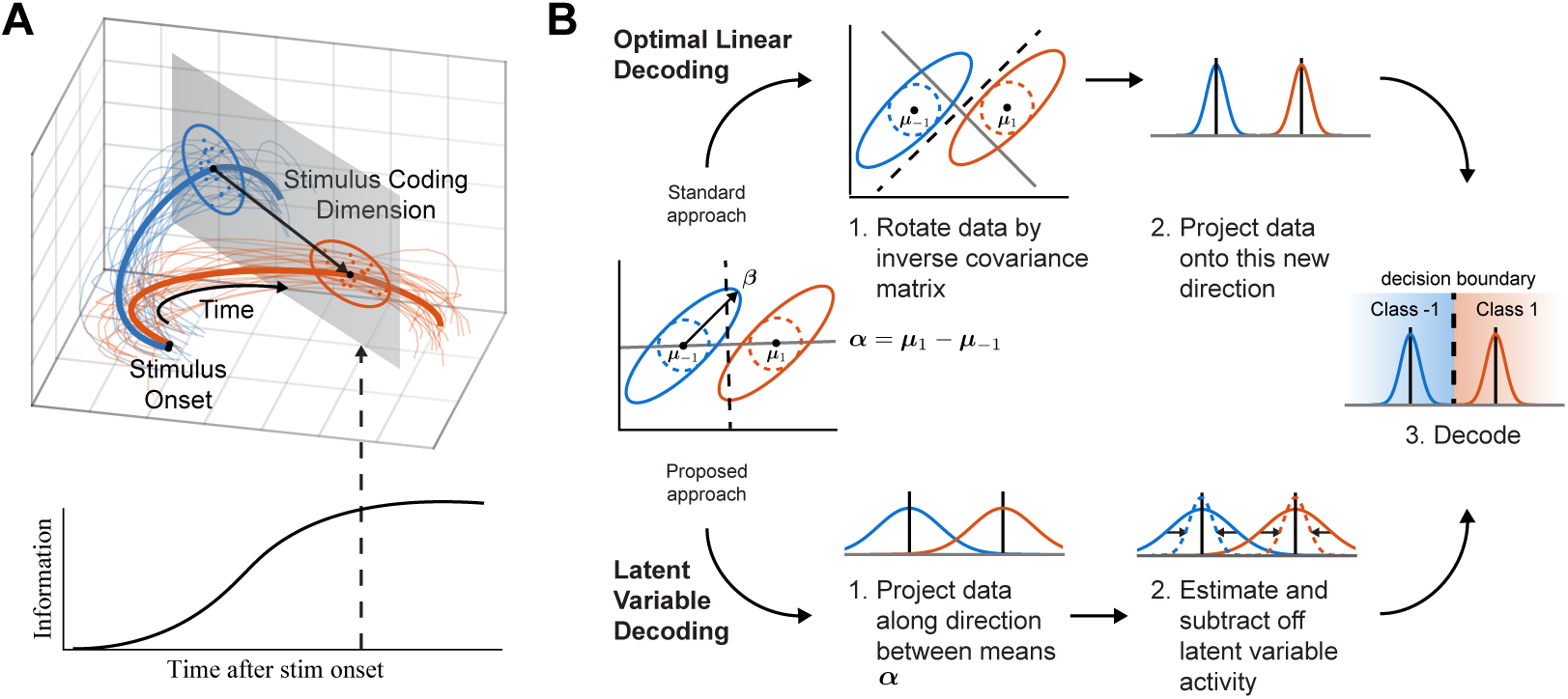
Using latent variables to decode stimulus identity from neural population activity. **A**: Illustration of the dynamics of high dimensional neural activity during a visually-guided saccade task, visualized using a latent variable model fit to trial-averaged activity. The trial-averaged activity at stimulus onset is marked by black dots, and evolves over time according to the saccade direction (leftward saccade, *bold blue line*; rightward saccade, *bold red line*). Activity from individual trials can also be projected into this space for visualization (*thin lines*). At a given point in time (*gray plane*), the saccade direction can be decoded from the neural activity by considering the position of the activity along the stimulus coding direction (*straight black arrow*). The more accurately the direction can be decoded from neural activity, the more information the population contains about the stimulus. **B**: Instead of using latent variables to visualize dynamics, the latent variable decoder exploits the covariance structure of the neural activity to remove variability that is shared among many neurons to improve decoding (*bottom*). This approach offers an alternative to optimal linear decoding (*top*), which accounts for the covariance structure by rotating the decoding direction by the inverse covariance matrix.

A full understanding of neural computation, however, requires understanding this single-trial activity. For example, trial-averaged trajectories reveal nothing about the differences between correct and error trials. A straight-forward way to visualize single-trial activity is to project it into the space defined by the trial-averaged data (Fig. 1A, *thin lines*), although, as noted above, this space will often fail to capture a large portion of the single-trial variability.

To help make the connection between noise correlations and latent variables, we consider neural activity at a single time point *t* during the experiment, and simplify the picture by only visualizing the neural activity in the two-dimensional plane that passes through the mean trajectories at time *t* (Fig. 1A, *gray plane*; Fig. 1B, *left*). Furthermore, we assume at time *t* the activity of neuron *n* on trial *i* is the sum of three terms (and drop the dependence on *t*): (1) the response to stimulus *s*_*i*_, with coupling strength *α*_*n*_; (2) the response to a latent variable *z*_*i*_ (described more below), which is independent of the stimulus, with coupling strength *β*_*n*_; and (3) a noise term 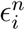, which is independent of both the stimulus and the latent variable:

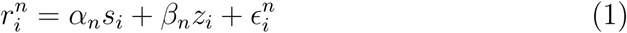

The latent variable *z*_*i*_ introduces trial-to-trial variability that is shared across the population, and while it might represent meaningful internal signals such as attention or arousal [26, 27], it could also arise due to internal structure within the network without clear behavioral relevance [28, 29]. The noise term 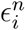, on the other hand, represents trial-to-trial variability that is private to each neuron. If we define the variance of *z*_*i*_ as 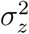 and the variance of 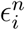 as 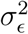, then the covariance matrix of the full population activity 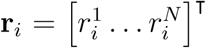, conditioned on the stimulus (an unscaled version of the noise correlation matrix), is given by

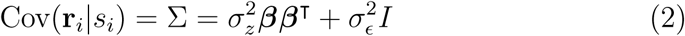

where ***β*** is the vector of all *β*_*n*_’s. The component due to the private noise term 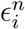 is a scaled version of the identity matrix (for simplicity; more generally it could be any diagonal matrix), and thus produces variability that is isotropic in space (Fig. 1B, *left, dashed circles*). The component due to the latent variable is a rank-1 matrix given by the outer product ***ββ***^T^, which highlights how the latent variable induces a low-dimensional structure on the noise covariance matrix. This latent variable component produces variability that is oriented along the direction ***β*** (Fig. 1, *left, solid ellipses*). This example generalizes to *K* latent variables, which would produce a rank-*K* component in the noise covariance matrix (and point along *K* different dimensions in the neural response space).

### 2.2 Decoding in the presence of latent variables

How does the presence of the latent variable affect the amount of information the neural responses contain about the identity of the stimulus on trial *i*? A straightforward approach for measuring linear information is to train a linear decoder to predict the stimulus using the neural activity (although see [30] for a direct estimation approach). Using the decoding approach, the optimal linear estimate of the stimulus *s*_*i*_ is defined as

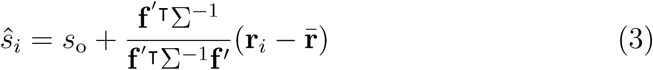

where *s*_o_ is the average stimulus, **f**′ is the vector of tuning curve derivatives with respect to *s*_*i*_, and 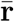 is the average population activity vector [31]. Linear Fisher information quantifies the accuracy of this estimate, and is defined as the inverse of the variance of *ŝ*_*i*_ [31].

Because the noise covariance matrix Σ plays a central role in the optimal linear decoder’s estimate of the stimulus, theoretical work has often addressed how decoding is affected by structure in this matrix that can arise in the neuroscience-related setting. These considerations include the impact of noise correlations that are related to the signal correlations [32], the relationship between diagonal and off-diagonal elements [33], and a noise component that points along the direction of individual neurons’ tuning curves [34]. However, no one has yet to our knowledge explicitly considered the consequences of a low-dimensional component arising from latent variables on decoding.

The optimal linear decoder takes the structure of Σ into account by multiplying ***α*** by Σ^−1^ to construct a decision boundary (Fig. 1B, *top*). This procedure rescales the space of neural responses to account for variances and cross-covariances among the different dimensions.

Our latent variable decoding framework proposes an alternative approach to the optimal linear decoder by assuming that the noise covariance matrix contains low-dimensional structure. We first project the full, high-dimensional neural activity onto the direction of ***α*** (Fig. 1B, *Latent Variable Decoding Step 1*), which in general is not the optimal decoding direction. Next, we form a trial-by-trial estimate of the variability in the direction of ***α***. It is possible to form this estimate because of the latent variable’s shared effect on the activity of the whole population; it would be impossible to form this estimate from the activity of any individual neuron, since its impact on neural activity is indistinguishable from the noise term 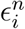 without the single-trial statistical power gained from simultaneously recorded neurons. Next, we take the estimate of this variability and subtract it from the projected population activity to reduce variability in the activity along this direction (Fig. 1B, *Latent Variable Decoding Step 2*), then finally decode the adjusted neural activity by comparing to a threshold value (Fig. 1B, *Latent Variable Decoding Step 3*).

Although this linear latent variable decoder cannot, by definition, outperform the optimal linear decoder in the limit of an infinite number of trials, we show in the following section that this decoder uses data more efficiently than other linear decoders, requiring fewer trials to extract the same amount of information. An additional feature of this framework is that it is not restricted to a linear method; indeed, using any nonlinear regression technique such as a neural network to estimate variability in the direction of ***α*** will result in a nonlinear latent variable decoding algorithm, which we also explore in the following sections.

### 2.3 Validating the LV decoder with simulated data

We demonstrate the performance of the latent variable (LV) decoder on simulated data, where it is possible to compare its performance to ground truth. Responses from 200 neurons were generated in a manner similar to Eq. 1, so that the same low-dimensional covariance matrix describes the variability around each of two mean responses (see Methods for simulation details). This data allows for an analytic expression of the linear Fisher information, and thus serves as a useful test case for our method. As a reminder, linear Fisher information provides an upper bound on the amount of information about the decoded variable that can be extracted from the population using linear operations.

To evaluate the performance of various decoders on this data we estimated Fisher information by calculating *d*′^2^ in the learned decoding direction (see Methods), and compared this to the true linear Fisher information (Eq. 15). *d*′^2^ is a quantity that must be estimated from data, and even the optimal linear estimator cannot extract the full linear Fisher information from limited data.

We tested performance of the LV decoder as a function of training trials (Fig. 2). Both the Linear and Nonlinear LV decoders extracted a large fraction of the true linear information using relatively few trials (Fig. 2A). Because this data does not contain nonlinear information, the Nonlinear LV decoder cannot perform better than the Linear LV decoder. We also show the performance of the “Difference of Means” decoder (Fig. 2A, *purple dashed line*), which uses the same decoding direction as the LV decoders, but does not estimate and subtract off shared variance. The gap in performance between the Difference of Means and LV decoders demonstrates the extent to which the LV decoders are able to account for shared variability that is detrimental to decoding (Fig. 2B). [See Fig. 3 for a comparison between the Linear LV decoder and other standard linear decoders on this simulated data.]

**Figure 2:**
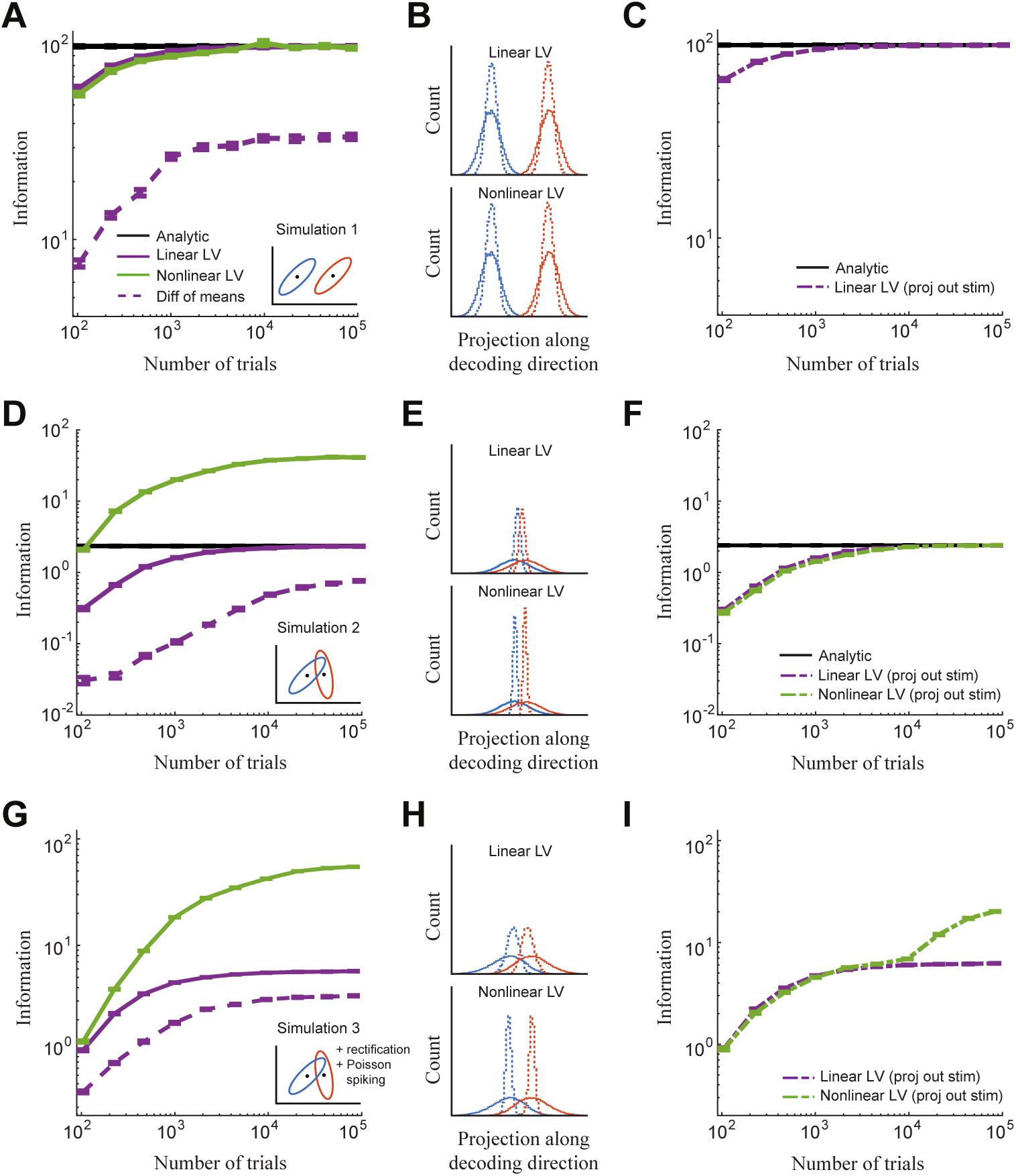
Linear and Nonlinear LV decoding performance on simulated data. **A**: Gaussian data is generated using the same covariance matrix for each class (*inset*), which only contains linear Fisher information (*black line*). Information measure is the *d*′^2^ discriminability index described in section 4.3, and error bars represent SEM on validation data over 25 simulated datasets. **B**: Histograms of the data projected onto the decoding direction for one dataset, colored by class, both before (*solid lines*) and after (*dashed lines*) subtracting off the predicted variability using the Linear LV (*top*) or Nonlinear LV (*bottom*) decoders. **C**: Decoder performance when projecting the stimulus dimension ***α*** out of the population activity before using it to infer variability. **D–F**: Same as *A-C*, but data is generated using a different covariance matrix for each class (see inset in *D*). **G–I**: Same as *D-F*, but the resulting values are rectified and passed through a Poisson spike generator to simulate spike count data.

**Figure 3:**
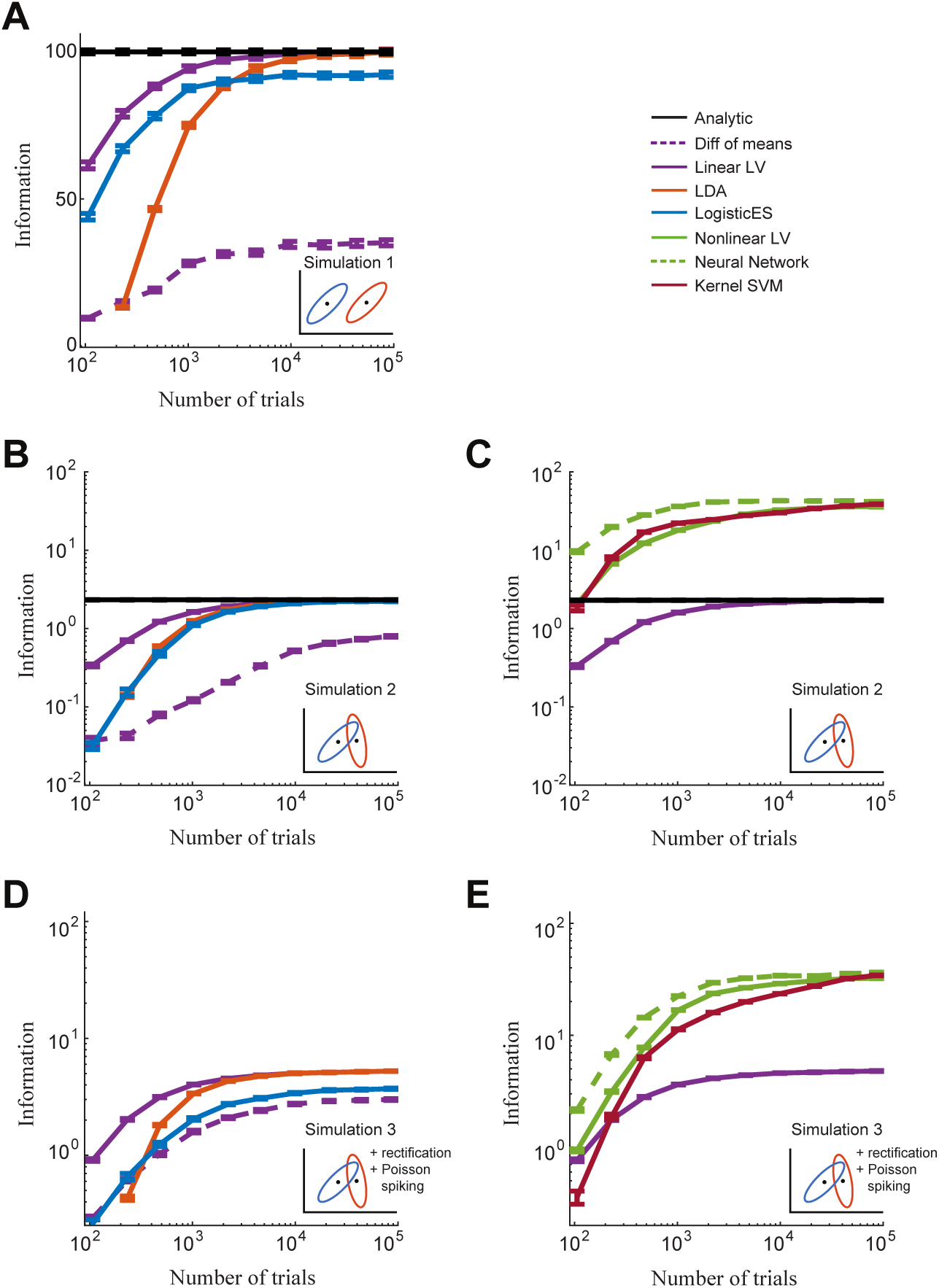
Linear and Nonlinear LV decoding performance on simulated data. **A**: Gaussian data with a single noise covariance matrix for both classes. **B, C**: Gaussian data with a different noise covariance matrix for each class; *B*: comparison of linear decoders; *C* : comparison of nonlinear decoders. **D, E**: Same as *B* and *C*, except resulting values are rectified and then passed through a Poisson spike generator to simulate spike count data. Details of the simulations are presented in the Methods.

To gain some intuition about how the neural networks might learn to predict the variability that is due to the latent variables, we analytically work out a simple example in the Appendix. This example suggests that the Linear LV decoder should project the high-dimensional neural activity onto a direction orthogonal to the stimulus coding dimension ***α*** to produce an estimate of the variability. To test this, we projected the ***α*** direction out of the data and retrained the Linear LV decoder, and found that its performance did not change (Fig. 2C).

The amount of information that a linear decoder can extract from data is bounded above by the linear Fisher information. However, a system with information that is not linearly decodable (for example, when the decision boundary is a curve rather than a straight line) would contain more information than just the linear Fisher information; in some cases, this nonlinear component of the full Fisher information can be substantially larger than the linear component [35].

One context where neural population activity contains nonlinear Fisher information is in the presence of stimulus-dependent noise correlations [36, 37]. We introduce this structure into the simulation by defining a different covariance direction for each stimulus context (Fig. 2D, inset), in which case the simulation contains nonlinear information [33], unlike the previous simulation, and thus serves as a natural extension for testing nonlinear decoders.

Figure 2D-F (analogous to Fig. 2A-C) demonstrates that the Nonlinear LV decoder can extract nonlinear Fisher information that is orders of magnitude larger than the linear Fisher information, and can do so with even a small number of trials. How does the decoding strategy learned by the Nonlinear LV decoder compare to that of the Linear LV decoder? We saw previously that the Linear LV decoder could estimate variability in a direction orthogonal to the stimulus coding direction (by projecting out the stimulus coding direction before decoding); when we performed the same experiment with the Nonlinear LV decoder, we found that its performance decreased substantially, becoming equivalent to that of the Linear LV decoder (Fig. 2F). This points to the ability of the Nonlinear LV decoder to extract nonlinear information by explicitly using activity in the direction of ***α***.

To test whether the LV decoders still perform well in a more realistic setting, we added additional statistical features of neural data to the simulation: we introduced stimulus-dependent noise correlations, followed by rectification and a Poisson spike generator to mimic the discrete nature of spiking data. These additional statistical features did not impair the performance of either LV decoder (Fig. 2G-I), demonstrating the ability of these techniques to generalize well to non-Gaussian data. Note that in this simulation the Nonlinear LV decoder can extract more information than the Linear LV decoder *without* using activity in the direction of ***α***, but only with a large number of training trials (Fig. 2I).

Finally, we compared the LV decoders with a range of other decoders, including linear discriminant analysis (LDA), logistic regression with early stopping regularization (LogisticES), kernel support vector machine (kernel SVM), and a two-layer neural network (Neural Network) (Fig. 3). In all three simulations the Linear LV decoder was able to achieve better performance than the other linear decoders (LDA and LogisticES) with a small number of training trials (Fig. 3A, B, D). The Nonlinear LV decoder, however, does not perform as well as a standard feed-forward neural network in the simulations that contain nonlinear Fisher Information (Fig. 3C, E). This suggests that while first estimating variability and then subtracting it out of the data is an effective approach for linear decoding, optimizing a cost function that is more directly related to classification accuracy is more effective in the nonlinear regime (at least with the simulations considered here). Additionally, the Nonlinear LV decoder performs equivalently to kernel SVM for simulation 2 (Fig. 3C), but attains better performance in the more realistic scenario of simulation 3 (Fig. 3E). [The same conclusions hold when using fraction correct rather than Fisher Information as a measure of decoder performance; results not shown.]

## 3 Discussion

We presented a novel method for decoding neural population activity, motivated by the observation that noise correlations in neural data can be effectively modeled using a small number of latent variables. As we showed, dimensionality reduction - which is common in neural population analyses (Fig. 1) - can be explicitly used by decoding methods to infer the variability along a chosen decoding direction and remove it, and thus eliminate its detrimental effect on decoding. While a linear latent variable (LV) decoder cannot outperform optimal linear decoding with infinite data, it is more efficient in its use of data than other common linear decoders for the range of simulations tested (Figs. 2 and 3). We also generalized this framework to nonlinear LV decoding, and demonstrated that it can extract nonlinear information approximately as well as other nonlinear LV decoding techniques (Figs. 2 and 3).

### 3.1 Decoder assumptions

The decoding framework presented here considers a binary classification problem rather than a continuous regression problem. We made this choice based on the rich literature that has developed around two-alternative forced choice tasks, which is the most natural setting for our LV decoding framework. Sani et al. [38] recently introduced a decoding technique that is similar in spirit to ours which considers the regression problem. Their technique finds a low-dimensional subspace of neural activity that is most predictive of behavior while considering other dimensions to contain non-relevant variability (with respect to the decoded behavior).

Our framework explicitly defines neural variability as resulting from LVs that are added to a mean stimulus response. Recent experimental results suggest that additive LVs may be present in cortical population activity [20,24], although other results suggest the LVs may be multiplicative [21, 23], or both [22, 25, 39]. Furthermore, our linear decoding framework only considers LVs that are not functionally targeted, i.e. a neuron’s coupling to the LVs is assumed independent of its stimulus tuning properties (although the Nonlinear LV decoder can in principle learn mappings from neural activity to LVs that are stimulus-dependent). Recent work by Haimerl et al. [40] proposed a decoding algorithm that considers neural responses which are functionally targeted by a multiplicative LV. The aim of their approach is focused on a flexible, biologically plausible decoder, while our approach is more focused on maximizing information extraction using as few training trials as possible. Nevertheless, our approach could be augmented by incorporating structure resulting from multiplicative LVs and functionally-targeted LVs (both additive and multiplicative) into the model of the noise covariance matrix.

### 3.2 The function of noise correlations

Variability of neural population activity along the stimulus coding direction (***α*** in Fig. 1B) cannot be distinguished from changes in the stimulus itself, and as such this form of variability is known as *information-limiting* or *differential* noise correlations [34]. Our decoding framework, although designed to reduce variability in the coding direction, does not remove information-limiting correlations (and indeed cannot, by definition). Instead, our decoder acts on variability that points in other directions of neural activity space (and therefore infers non-information-limiting noise correlations), and removes the projection of this variability into the coding direction (the projection of ***β*** onto ***α*** in Fig. 1B).

Although the LV decoder was not designed with strict biological plausibility in mind, it provides a useful framework for considering how the brain might deal with this type of variability. The sensitivity of noise correlations to behavioral context [41, 42], attention [27, 43, 44] and perceptual learning [27, 45] suggest that they are at least partially the product of top-down, feedback processes in sensory cortex. However, these types of correlations can reduce the amount of information available in a neural population of finite size [33] (though not limit the information as the population becomes infinitely large [34]). What then is the functional role of these correlations?

Theoretical work has shown how inducing particular patterns of noise correlations improves information transmission between different brain regions [46]. Another line of theoretical inquiry implicates correlated variability in the representation of prior information in a Bayesian framework of sensory integration [26, 47]. Here we show that the goals of robust information propagation or perceptual inference (where noise correlations can help) and accurate decoding (where noise correlations can hurt) are not necessarily at odds with one another. We hypothesize that if activity in a brain region is corrupted by latent-variable-induced noise correlations, a downstream decoder of that activity need not be negatively impacted if it has access to the top-down signal inducing the correlations (a possibility also addressed by [26, 40, 42]). Decision-making areas could then conceivably integrate sensory information with these top-down signals [47, 48], or use them for flexible, task-dependent information routing [40].

Our results are also consistent with a recent study demonstrating that the inclusion of activity from untuned neurons can increase the performance of decoders [49]. In our framework, these neurons might not be tuned to the particular task, but they can still carry information about the signals that give rise to correlated variability. Including these neurons in a decoder can then lead to more accurate estimation of the trial-to-trial variability, which in turn will improve the performance of the decoder.

### 3.3 Application to real data

The application of the LV decoders to experimental data is straightforward. However, our simulations encode very simple assumptions about sources of variability, and it is not clear how well the LV decoders we developed here will generalize to other commonly encountered sources of variability in neural data. These sources could be introduced by the recording modality, such as electrode drift in electrophysiology data, or bleaching in calcium imaging data. Sources of variability could also be introduced by uncontrolled fluctuations in the state of the animal, such as changing levels of arousal, attention, or locomotion. These sources of variability (as well as many others not considered here) most likely have more complex structure than the basic Gaussian noise terms built into our simulations. Comparing the LV decoders to other common decoding algorithms in these settings is a clear avenue for future work.

## 4 Methods

### 4.1 The LV decoder

#### Training the LV decoder

For a population of *N* neurons recorded during *T* trials, we define *R* = [**r**_1_ … **r**_*T*_]^T^ ∈ ℝ^*T*×*N*^ to be the matrix of spike counts and **y** = [*y*_1_ … *y*_*T*_]^T^ ∈ {±1}^*T*^ to be a binary vector that indicates the stimulus identity *s* ∈ {±1} on each trial *i* ∈ {1, …, *T*}. The two stimuli could be, for example, gratings with different orientations in a visual paradigm, or tones with different frequencies in an auditory paradigm. The following steps are illustrated in Fig. 4.

**Figure 4:**
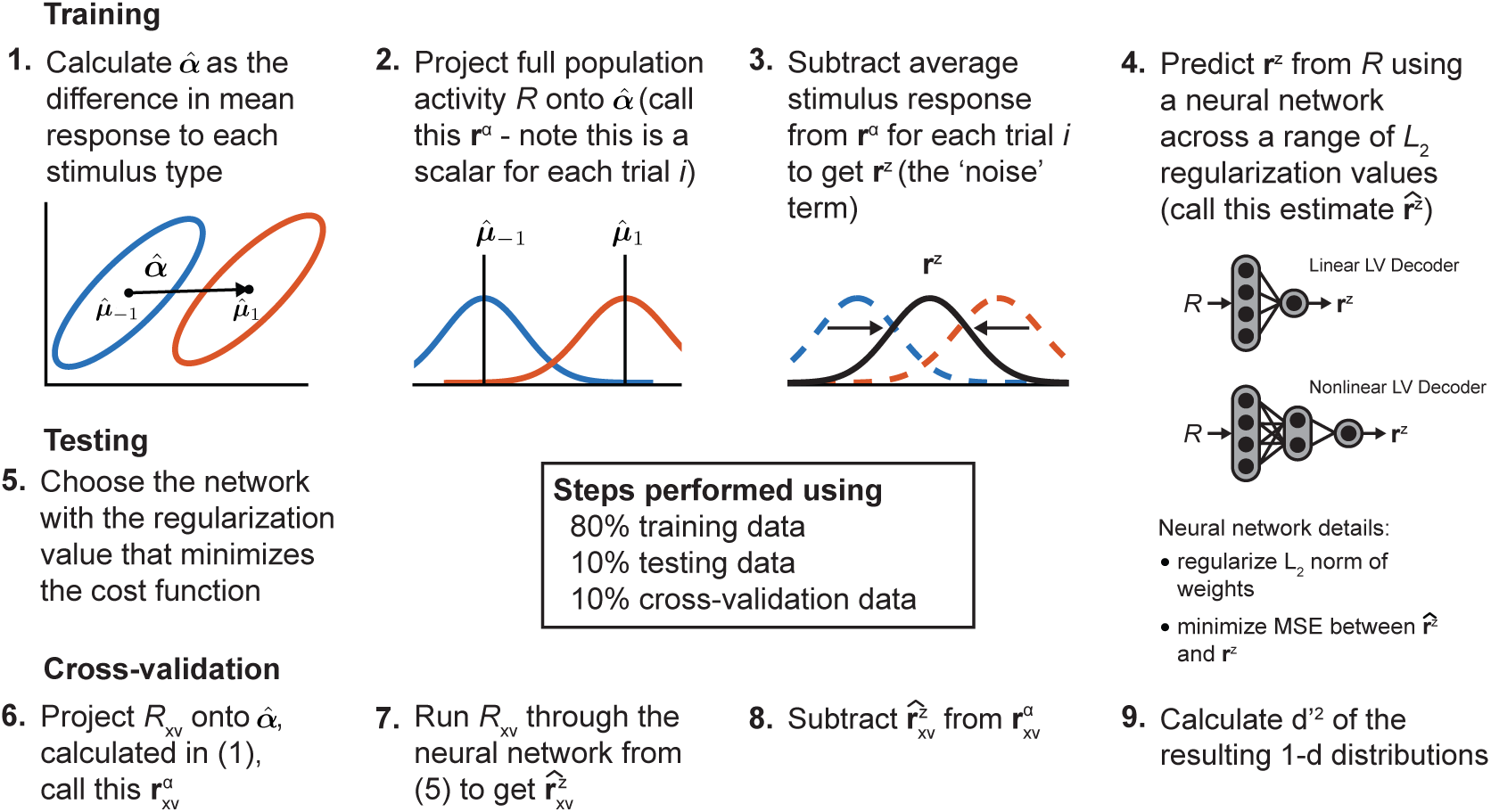
Outline of LV decoding algorithm.

**Step 1:** We first estimate the mean stimulus responses 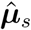 as

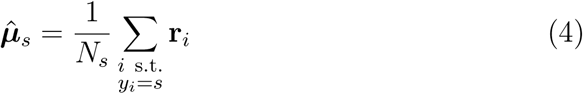

where *N*_*s*_ is the number of stimuli from class *s*. We then estimate the stimulus coding direction 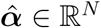 that points between the two mean stimulus responses as 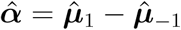.

**Step 2:** Next, we project *R* onto 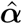 to reduce the high-dimensional neural activity into a single dimension,

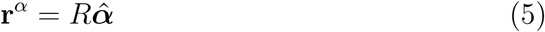

which is now a scalar value for each trial *i*^1^.

**Step 3:** We next estimate variability in the stimulus coding direction that is not stimulus-driven. To do so, for each trial *i* we subtract the appropriate mean stimulus response from the projection along 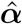 and denote this new quantity **r**^*z*^ :

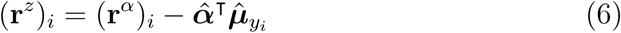

where (**x**)_*i*_ denotes the *i*th component of a vector **x**. The extent to which **r**^*z*^ contains variability that is shared across many neurons (due to latent variables) determines the efficacy of the LV decoder.

**Step 4:** We estimate this shared component of variability with the full population response *R* by learning a mapping *f*_*θ*_ : ℝ^*N*^ → ℝ parametrized by *θ* using a neural network (see *Neural network details* below), so that

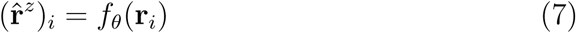

#### Evaluating the LV decoder

Once *f*_*θ*_ has been learned, we can evaluate the performance of the LV decoder. As an example, we will use *R*_xv_ to represent the cross-validation data. *R*_xv_ is first projected onto 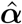 (learned from the training data) to get 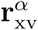 (*Step 6*). The activity **r**_xv_ from each trial is then run through the function *f*_*θ*_ to produce one component of the vector 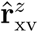 (*Step 7*). Finally, the variance-reduced activity is given by (*Step 8*)

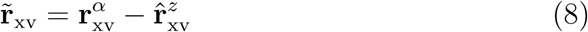

To calculate the classification of each trial *i*, the corresponding value from 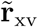 is compared with 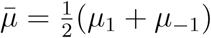; values larger than this threshold are classified as stimulus 1, and all others are classified as stimulus −1 (*Step 8*).

#### Neural network details

Here we describe details of the neural network that is used to estimate variability in the stimulus coding direction. Any technique that can learn a mapping from ℝ^*N*^ to ℝ is suitable in principle, but we restrict our explorations to a standard neural network for simplicity. The neural network takes *R* as input and produces an estimate 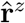 of **r**^*z*^. Parameters of the network *θ* are learned by minimizing the mean square error (MSE) between 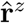 and **r**^*z*^. *L*_2_ regularization is included to prevent overfitting to the training data [50], so that the penalized cost function *C* is defined as:

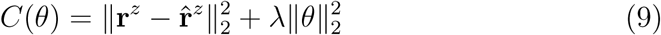

where 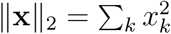 is the *L*_2_ norm of a vector **x** and *λ* is a hyperparameter that controls the magnitude of the regularization term. In practice, we fit the LV decoders using 10 different values of *λ* logarithmically spaced between 1*e*-4 and 1*e*1, and choose the value that results in the smallest cost function when evaluated on the testing data (*Step 5*). The cost function is optimized using an L-BFGS routine [51].

The Linear LV decoder requires a linear mapping from ℝ^*N*^ to ℝ, and therefore uses a neural network with just an input layer and an output layer. With the *L*_2_ regularization, this network is equivalent to regularized linear regression, or “ridge regression” [50]. The Nonlinear LV decoders use a neural network with a single hidden layer composed of 15 rectified linear units (ReLUs), which we found to work well for all simulated datasets. We explored different numbers of hidden units and hidden layers, but did not perform an exhaustive hyperparameter search.

#### Projecting out the stimulus coding dimension

To test the extent to which the LV decoders require information contained in the stimulus coding direction ***α*** (Fig. 2C, F, I), we projected this dimension out of the population activity *R* before using it to predict variability in the same dimension ***α***, and we denote the resulting activity by 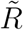. The stimulus coding dimension was calculated after subsampling trials, and was only calculated using training data. Training the decoder then amounted to replacing *R* in Eq. 5 with 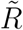. To evaluate the decoder, the same ***α*** was projected out of the testing/cross-validation data, and all other steps in the *Evaluating the LV decoder* section remain the same.

### 4.2 Training of standard decoders

#### Difference of means (DoM)

The mean response to each stimulus was calculated; the difference in mean responses ***α*** defined the discriminant line for the DoM decoder, and the mean of the mean responses defined the threshold. For each trial, neural activity was projected onto the discriminant line and compared to the threshold value to determine its classification.

#### Linear discriminant analysis (LDA)

LDA was performed using the `fitcdiscr` function in MATLAB, with the ‘*DiscrimType*’ option set to ‘*linear* ‘ so that a single pooled covariance matrix was estimated from the data. The ‘*Gamma*’ option was set to 0, so that the estimated covariance matrix was not regularized with an additional diagonal matrix. This choice limited the use of LDA to settings where the number of trials was larger than the number of neurons.

#### Logistic regression with early stopping (LogisticES)

Logistic regression models were fit by minimizing the mean square error between class labels *y* ∈ {0, 1} and predicted class labels given by

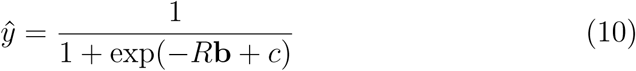

where *R* is the matrix of neural responses, **b** is the vector of learned decoder weights and *c* is a learned bias term. The negative log-likelihood of the testing data was evaluated on each iteration, and model fitting terminated once the negative log-likelihood began to increase or the algorithm reached 1000 iterations [52].

#### Neural network decoder

Neural networks were used as an additional nonlinear decoding algorithm. The networks were trained to take in neural population activity and predict the stimulus class (±1); parameters were learned by minimizing penalized MSE between true and predicted class using the L-BFGS routine (*L*_2_ regularization was applied to the weights using the same range as the LV decoders). The network architecture matched that of the Nonlinear LV decoder - a single hidden layer comprised of 15 ReLU units.

#### Kernel support vector machine (Kernel SVM)

Kernel SVMs were fit using the `fitcsvm` function in MATLAB with the ‘*KernelFunction*’ option set to ‘*rbf* ‘ to use radial basis function kernels. Radial basis functions are unnormalized Gaussians, and the scale of these functions relative to the data is important for kernel SVM performance. MATLAB provides another option ‘*KernelScale*’ that scales the data (rather than the kernel); to fit this hyperparameter, we fit kernel SVMs using 10 different values of the scale parameter logarithmically spaced between 1*e*-3 and 1*e*3, and chose the scale that resulted in the largest number of correctly classified trials when evaluated on the testing data (see *Evaluating decoder performance* below).

### 4.3 Evaluating decoder performance

#### Subsampling trials and cross-validation

A main goal of this study was to understand how the performance of different decoders scaled with the number of trials. To do so we first removed 10000 trials from the data as validation trials, which were not used for training or hyperparameter selection. Then for each dataset size (*K* = 100 to *K* = 90000 trials), we randomly sampled *K* trials from the dataset, then randomly split these trials into five folds - four for training and one for hyperparameter selection (e.g. *L*_2_ regularization for LV and Neural Network decoders, early stopping for LogisticES decoders, etc.). The best model, found using the testing data, was then evaluated on the held-out validation trials, and these are the values reported in Figs. 2 and 3.

### Quantifying decoder performance with *d*′

The simplest measure for quantifying decoder performance is the fraction of correctly classified trials. For LDA and kernel SVM, the predicted classification for each trial was obtained using the `predict` function in MATLAB. The DoM, LV, and Neural Network decoders explicitly define a threshold, and a trial is classified based on comparing the projection of the data along the learned discriminant line to the threshold. For LogisticES, the predicted class label *ŷ* (Eq. 10), a continuous quantity between 0 and 1, was turned into a binary classification by using 0.5 as a threshold.

However, in this work, our concern was not in the fraction of correctly classified trials, but rather in the total amount of information the neural population contains about the decoded variable. For example, when classes are fully separable, fraction correct is unable to distinguish between a decoder with high variance and one with low variance (see, for example, Fig. 2B). So instead of reporting fraction correct we instead use the more general *linear Fisher information* measure to quantify decoder performance.

Linear Fisher information measures the inverse of the variance of a decoder’s prediction of the stimulus, and therefore decoders with smaller variance in their predictions will contain more information. We estimate linear Fisher information using two different computations of the *d*′ measure^2^: 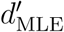, which is most appropriate when the simulation noise is Gaussian, and useful when the data are nearly or completely separable (simulation 1); and 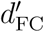, which is more appropriate when the simulation noise is non-Gaussian and the data are far from separable (simulations 2 and 3), and/or the decoding algorithm does not project the data along a discriminant line (such as kernel SVM).

#### Computing 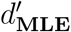

To compute the 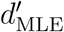 measure, the full-dimensional population activity is first projected onto the discriminant line (which precludes the use of this measure with kernel SVM, which does not estimate discriminant lines). In our simulated data the resulting one-dimensional projection for each class is well-described by a Gaussian distribution (e.g. Fig. 2B). The mean and variance of these distributions are fit for each class using the maximum likelihood estimates (MLE). Then, the fraction of correctly classified trials for each class (denoted as the accuracy *A*), in the limit of infinite data, is estimated by using the error function of the Gaussian defined by these MLE parameter values.

For example, if the mean of class −1 is located to the left of the threshold, the fraction of correctly classified trials from this class is given by the area under the curve between the threshold and negative infinity, and is denoted by *A*_−1_ (*A*_1_ is defined analogously for the other class). The overall fraction of correctly classified trials is then estimated as 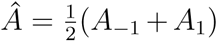, and this value can be converted to the *d*′ measure using the inverse of the complimentary error function *H* [33]:

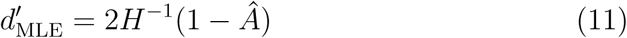

We use this computation for *d*′ in Simulation 1 (see Table 1), where classes are fully separable and projections onto the discriminant line are guaranteed to have a Gaussian distribution.

**Table 1:**
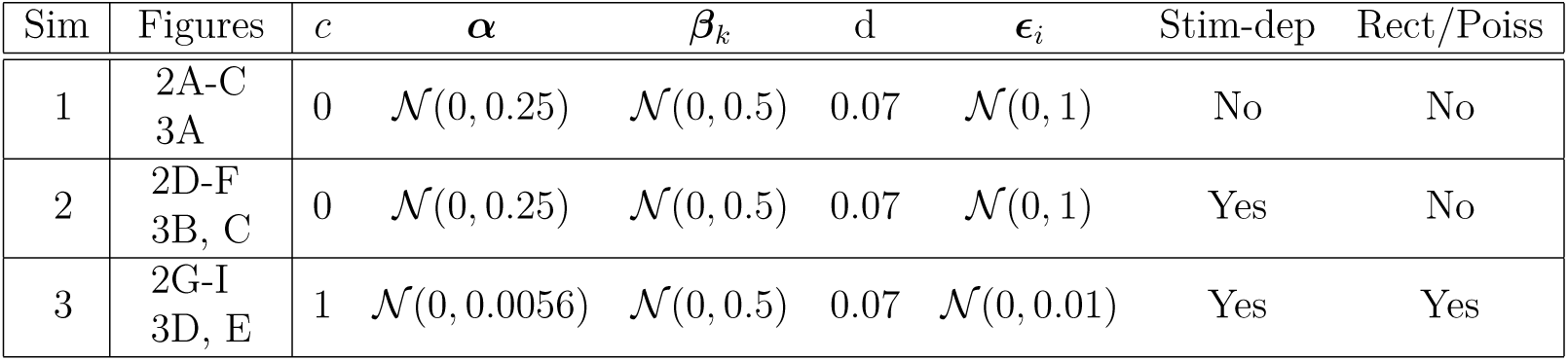
Simulated data details. The performance of various decoders evaluated on these simulated datasets is shown in Figs. 2 and 3. All datasets were generated using *N* = 200 neurons, *K* = 10 latent variables and *T* = 100000 trials.

#### Computing 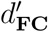

A drawback to 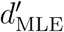 is that it cannot be used to evaluate decoding algorithms such as kernel SVM, which do not linearly project the data along a discriminant line. An alternative way to compute *d*′ (when classes are not fully separable) is to use the fraction of correctly classified trials, or accuracy *A* (*not* in the limit of infinite data, as before). As before, this value can be converted to *d*′ :

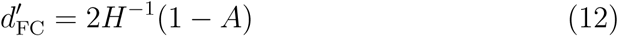

where ‘FC’ denotes ‘fraction correct’. We use this computation for *d*′ in Simulations 2 and 3 (see Table 1), where classes are not fully separable and projections onto the discriminant line are not guaranteed to have a Gaussian distribution.

#### Equivalence of linear Fisher information and *d*′^2^

Throughout this paper, we refer to *d*′^2^ as ‘Information’. To justify this equivalence, we show here that yet another definition of *d*′ is equivalent to linear Fisher information when calculated along the optimal coding direction and squared. Although the three values of *d*′ considered here differ in their computation, under the assumption of Gaussianity they become equivalent in the limit of infinite data.

We now consider the classic definition of *d*′ [53], which was originally introduced in the signal detection literature as a measure of the signal-to-noise ratio (SNR):

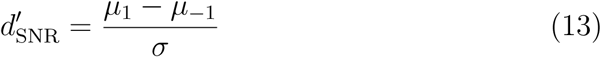

where *µ*_*i*_ is the mean of the *i*th one-dimensional response distribution and *σ* is the standard deviation, which we take to be the same for both distributions. If we now consider the response **r** of a population of neurons, with a stimulus-conditioned covariance matrix given by Cov(**r**|*s*) = Σ, the definition of linear Fisher information in this context becomes

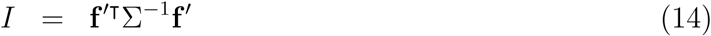

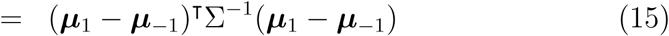

We now calculate 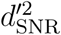 of the response distributions after they have been projected along the optimal decoding direction 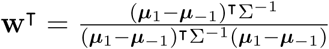 [31], and show that this is equivalent to the expression for linear Fisher information in Eq. 15.

The means of the response distributions along this dimension, denoted by 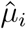, are 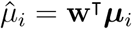 and the variance along this dimension (which we again assume is the same for both response distributions), denoted by 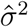, is

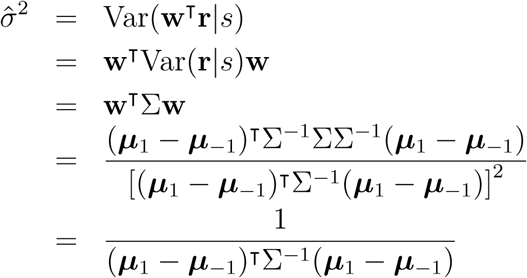

and thus

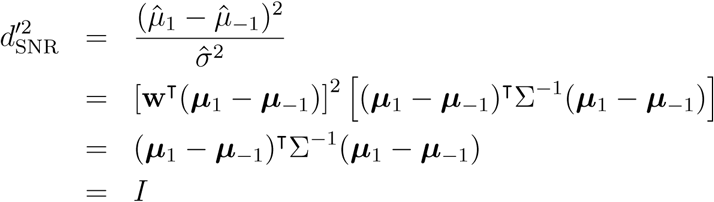

### 4.4 Simulation details

We tested all decoders on a variety of simulated datasets. For all simulations, we generated the responses of *N* neurons over *T* trials, where the population response **r**_*i*_ on trial *i* was generated as a sum of five terms: (1) a bias; (2) the stimulus *s*_*i*_ ∈ {±1}, coupled to the population via ***α***; (3) a collection of *K* latent variables 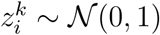 coupled to the population via ***β***; (4) a (*K* + 1)^st^ latent variable 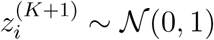 that points in the coding direction ***α*** with strength *d*, to explicitly introduce information-limiting noise correlations [34]; and (5) and a noise term *ϵ*_*i*_:

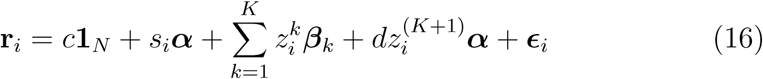

where **1**_*N*_ is a vector of *N* 1s. We assume that all statistical quantities in Eq. 16 are independent of each other. Data generated in this way results in a single noise covariance matrix that is independent of the stimulus identity:

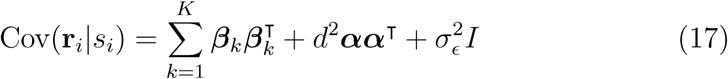

In this setting, linear discriminant analysis is equivalent to the optimal linear decoder, and the population response only contains linear information [33]. We introduced nonlinear information into the population via stimulus-dependent noise covariance matrices, which requires a separate, independent set of latent variable coupling vectors 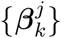 for each stimulus value *j*, so that

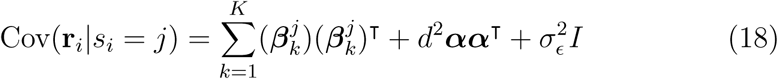

To generate data more closely resembling neural activity, for some analyses we rectified the values of **r**_*i*_, and the resulting non-negative values were used as rate parameters for independent Poisson processes to produce spiking activity. Details of each simulation are shown in Table 1. For each row of the table (corresponding to each row of the indicated figures) we randomly sampled 25 datasets; errorbars in the figures show SEM over the datasets.

## Acknowledgements

This work was supported by the NSF IIS-1350990 and NIH R21 EY025403-02 (MRW and DAB) and NIH ZIA MH002928-01 (BA).

## Appendix: Single latent variable example

This appendix provides a deeper analysis of the single latent variable example introduced in sections 2.1 and 2.2. To restate the problem formulation, the population firing rate vector **r**_*i*_ ∈ ℝ^*N*^ on trial *i* is the sum of three terms: (1) the stimulus *s*_*i*_ ∈ {±1}, coupled to the population via ***α***; (2) a latent variable 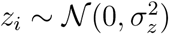 coupled to the population via ***β***; (3) and a noise term 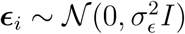:

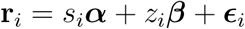

We assume that *z*_*i*_ and ***ϵ***_*i*_ are independent, so that Cov(*z*_*i*_, ***ϵ***_*i*_) = 0. To facilitate the derivations below, we make the further assumptions that ***α*** and ***β*** are unit vectors (more generally, the magnitude of each vector can be absorbed into the scalars *s*_*i*_ and *z*_*i*_), and that ***α*** and ***β*** are known. The covariance matrix of population activity, conditioned on *s*_*i*_, is given by

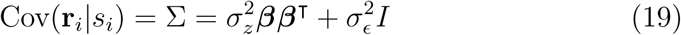

In the remainder of this appendix we derive an analytic “latent variable” estimator for *s*_*i*_ under these specific assumptions and examine its statistical properties in relation to the optimal linear estimator.

### A latent variable estimator for *s*_*i*_

The optimal linear estimator for *s*_*i*_, denoted by 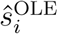, is

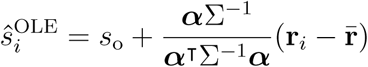

where *s*_o_ is the average stimulus value (0 in this case) and 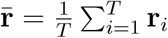 [31]. We propose to exploit our knowledge of the structure of Σ in Eq. 19 to derive a different estimator for *s*_*i*_. We will first infer the activity of the latent variable *z*_*i*_ in the direction of ***α***, then remove this component from **r**_*i*_ before decoding in the direction of ***α***.

We can infer the latent variable *z*_*i*_ by projecting the response vector onto ***α***_⊥_, the component of ***β*** that is orthogonal to ***α***:

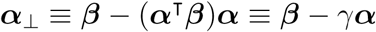

so that *γ* corresponds to the cosine of the angle between ***α*** and ***β***. Then the projection of the response vector along ***α***_⊥_ becomes

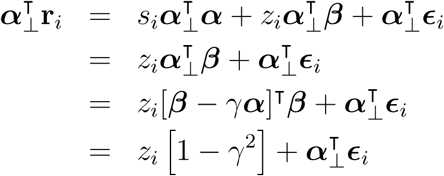

Rearranging,

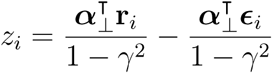

so that

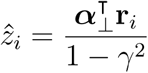

is an unbiased estimator for *z*_*i*_, and 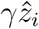 is an unbiased estimator for the projection of the latent variable term *z*_*i*_***β*** along the ***α*** direction.

To arrive at the latent-variable-adjusted estimate of the stimulus, *ŝ*^LVE^, we simply project the population activity along the direction of ***α*** and subtract the estimate of the latent variable term in that direction:

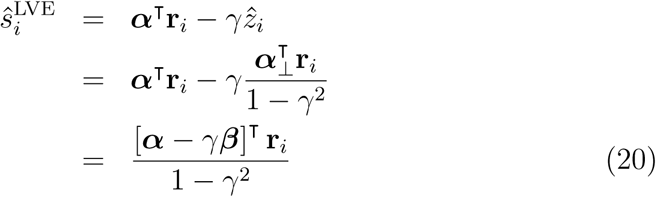

This estimate of the stimulus depends on the angle between ***α*** and ***β*** through *γ*, and it is instructive to note the two extreme cases. First, when ***α*** and ***β*** are parallel, *γ* = 1 and there is no solution, because *z*_*i*_ cannot be disambiguated from the stimulus (in this situation the induced correlations would be *information-limiting* noise correlations [34]). Second, when ***α*** and ***β*** are orthogonal, *γ* = 0 and the latent variable is not detrimental to decoding along ***α***, so that the estimate of the stimulus reduces to *ŝ*_*i*_ = ***α***^T^**r**_*i*_.

### Linear Fisher information for 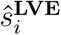

Given the estimate 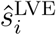 in Eq. 20, we can calculate its linear Fisher information as the inverse of the variance of 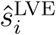, which is given by

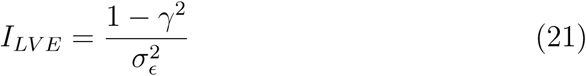

How does this compare to the linear Fisher information of the optimal linear estimator 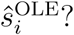 By substituting Eq. 19 into the standard result that *I*_OLE_ = ***α***^**T**^Σ^−1^***α***,

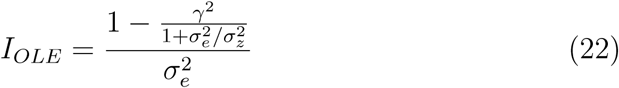

Again, we note the two extreme cases. When *γ* = 1, *I*_LVE_ = 0 because the latent variable is pointing in the direction of the stimulus, but *I*_OLE_ is greater than zero. This illustrates an important case in which 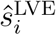 is far from optimal. When *γ* = 0, however, *I*_LVE_ and *I*_OLE_ are equivalent. Results from intermediate values of *γ* are shown in Fig. 5.

**Figure 5:**
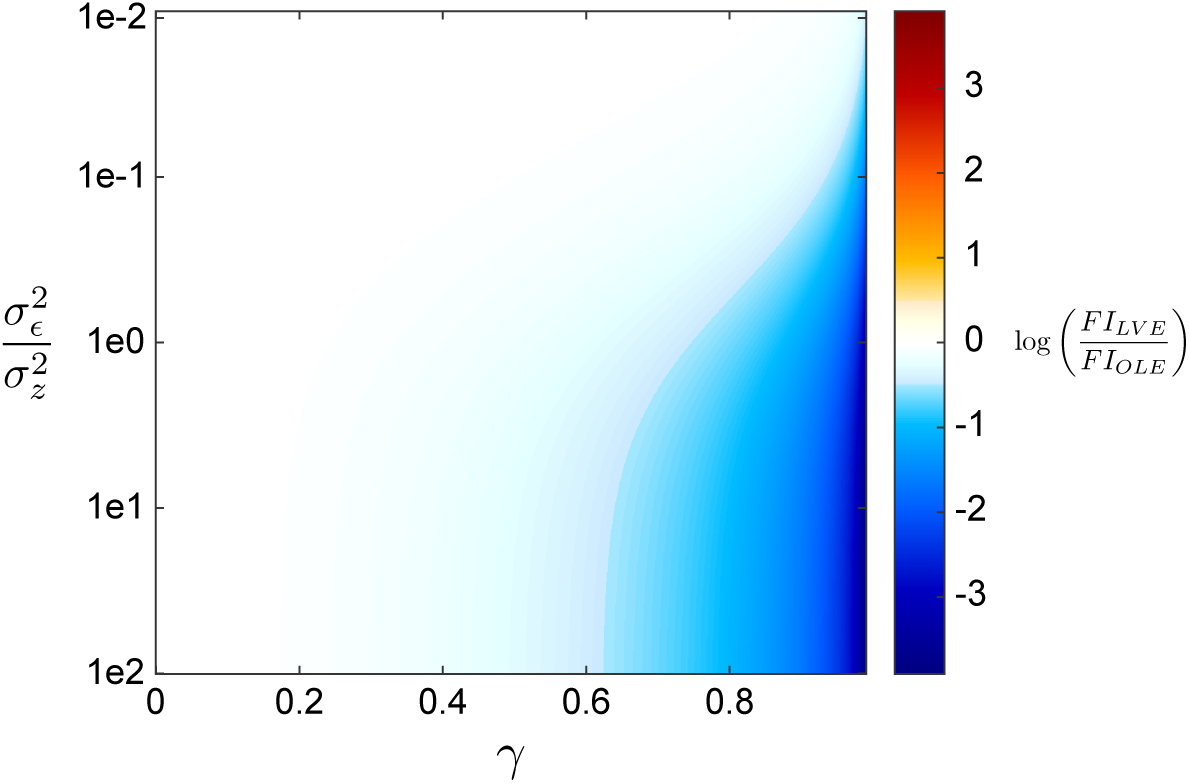
Comparison of estimators for the single latent variable model. Color indicates the logarithm of the ratio of the linear Fisher information for the latent variable estimator (*I*_LVE_, Eq. 21) and the optimal linear estimator (*I*_OLE_, Eq. 22). This value is plotted as a function of the ratio of the variances of the noise (*σ*_*ϵ*_) and the latent variable (*σ*_*z*_), and *γ*, the cosine of the angle between ***α*** and ***β***.

Why does *I*_LVE_ → 0 as *γ* → 1? This behavior is easier to understand by considering the variance of the estimate 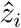, which is given by

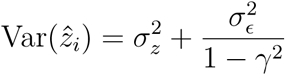

The variance of 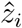 is equal to the variance of *z* plus a term that depends on *γ*. When ***α*** and ***β*** are orthogonal (*γ* = 0), this second term becomes equal to 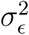, the variance of the noise. As ***α*** and ***β*** become more aligned (*γ* → 1), the variance of 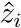 blows up and drives *I*_LVE_ to zero.

The LV decoder cannot, by definition, extract more information from population responses than the optimal linear decoder. However, this single latent variable example demonstrates that there are a wide range of parameter settings for which the LV decoder performs close to optimal. Importantly, this analysis only considers the behavior of these estimators in the limit of infinite data, and does not consider how efficiently these estimators use finite amounts of data. In practice (i.e. with a limited number of trials), the LV decoder is able to more efficiently extract information than other linear decoders (see Fig. 3).

## Footnotes

1 For reference, decoding **r**^*α*^ is referred to as the “Difference of Means” decoder used throughout the text. This decoder generally has worse performance than the LV decoder, even though both use the same decoding direction. Their difference demonstrates the extent to which variability in the direction of **r**^*α*^ is both (1) detrimental to decoding, and (2) shared across multiple neurons, and hence inferrable from the data.

2 see the section *Equivalence of linear Fisher information and d*′^2^ below for a non-rigorous proof demonstrating the equivalence of these two quantities under simple assumptions.

